# Short Tandem Repeats Information in TCGA is Statistically Biased by Amplification

**DOI:** 10.1101/518878

**Authors:** Siddharth Jain, Bijan Mazaheri, Netanel Raviv, Jehoshua Bruck

**Affiliations:** California Institute of Technology, Electrical Engineering, Pasadena, 91125, California, USA; California Institute of Technology, Computing and Mathematical Sciences, Pasadena, 91125, California, USA

## Abstract

The current paradigm in data science is based on the belief that given sufficient amounts of data, classifiers are likely to uncover the distinction between true and false hypotheses. In particular, the abundance of genomic data creates opportunities for discovering disease risk associations and help in screening and treatment. However, working with large amounts of data is statistically beneficial only if the data is statistically unbiased. Here we demonstrate that amplification methods of DNA samples in TCGA have a substantial effect on short tandem repeat (STR) information. In particular, we design a classifier that uses the STR information and can distinguish between samples that have an analyte code D and an analyte code W. This artificial bias might be detrimental to data driven approaches, and might undermine the conclusions based on past and future genome wide studies.

## Introduction

Genomic studies of complex diseases, commonly referred to as *Genome Wide Association Studies* (GWAS)^6^, flooded the literature soon after the first human genome was sequenced in the early 2000’s^8^. These studies aimed to map thousands of small risk variants that collectively affected the occurrence of the complex disease in question. The variants are detected by comparing the DNA of healthy individuals against sick individuals. As each complex disease was speculated to be caused by a large number of small risk variants, massive amount of genomic data was needed. This became the primary incentive for projects that collected massive amounts of genomic data, such as the 1000Genome Project^1^, The Cancer Genome Atlas (TCGA)^21^, and UK Biobank^17^, to name a few.

In addition, recent computational advances in machine learning (ML) tools, such as decision trees and neural networks, have ignited a data driven learning approach to tackle the complex associations between variants and their connection to disease risk. The DNA samples used in these studies undergo different amplification techniques and are derived from many different sources. These differences potentially contribute distinct noise patterns, specific to a given source or technique^4^, thus leading to biased data that could drastically alter the outputs of sensitive ML methods. We demonstrate the existence of such bias in TCGA and the vulnerability of ML methods to this bias.

## Results

To examine this bias, we study the effect of amplification techniques on whole exome sequencing (WXS) data on TCGA. This data comprises more than 10,000 exomes of both tumor and normal type (blood-derived or primary tissue) for 33 different cancers, which have been obtained from different sources and have undergone different amplification techniques. In particular, we compare WXS BAM files labeled with the analyte code “D” and files with the analyte code “W”. Samples with “W” code have undergone whole genome amplification (WGA) using Repli-G (Qiagen) technology that uses Multiple Displacement Amplification (MDA).

By merely using the copy number and number of point mutations in short tandem repeat regions of normal DNA, we are able to distinguish “D” and “W” category files of the same cancer with high accuracy (Experiment 1: see Figure 1). We then show that this signal contributes to misclassification when classifying DNA by cancer type (Experiment 2: see Figure 2). We describe the details of these experiments below.

**Figure 1.**
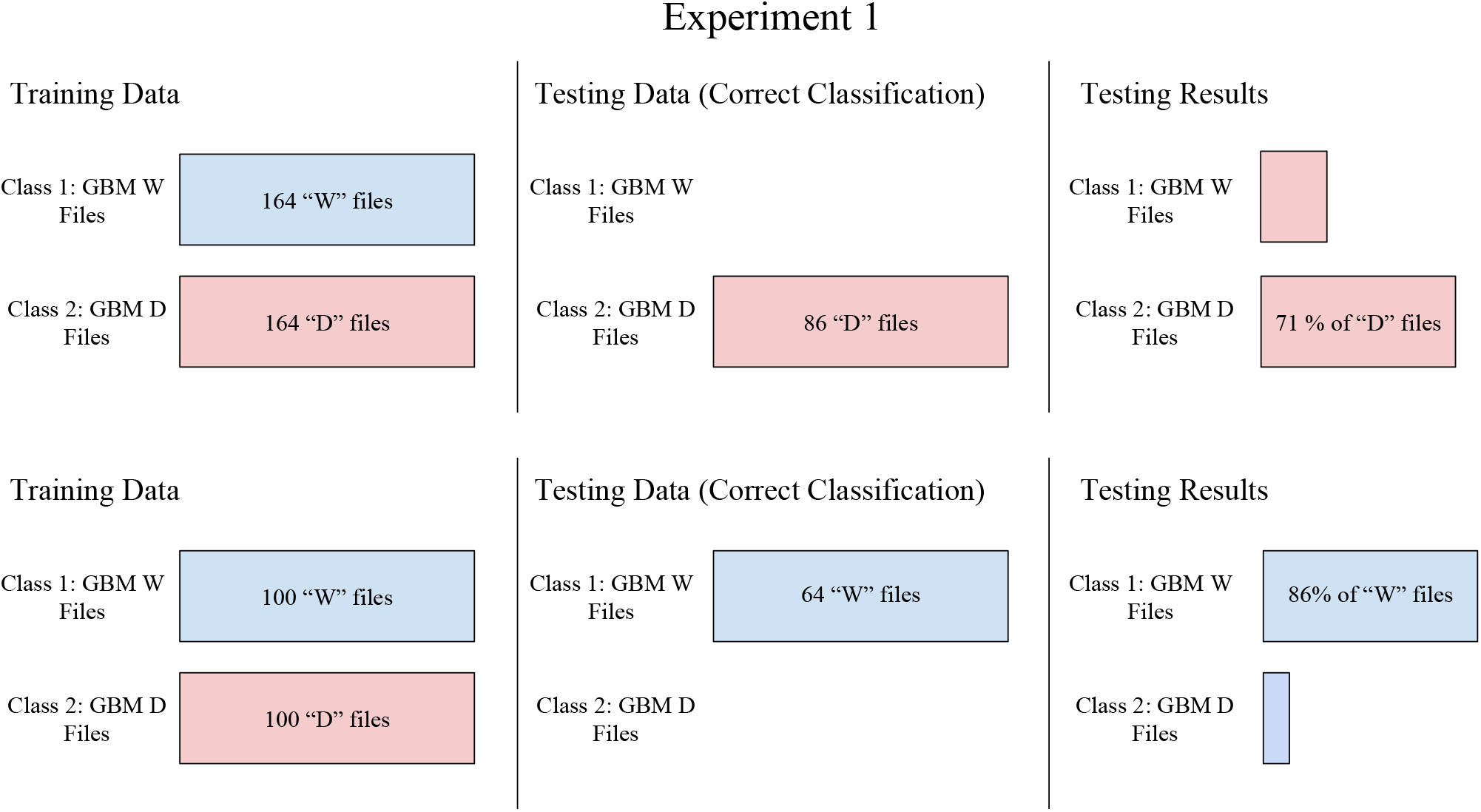
An illustration of experiment 1. Here, the placement of bars represent the labeling of the classes. Throughout our diagrams we color “D” type files red and “W” type files blue. The first column shows the data that our classifier was trained on. The second column shows where a perfect classifier would put the data, and the third column shows how our classifier labeled that data. Here we see that, within the cancer class of GBM, we are able to train a fairly accurate classifier for “D” and “W” files.

**Figure 2.**
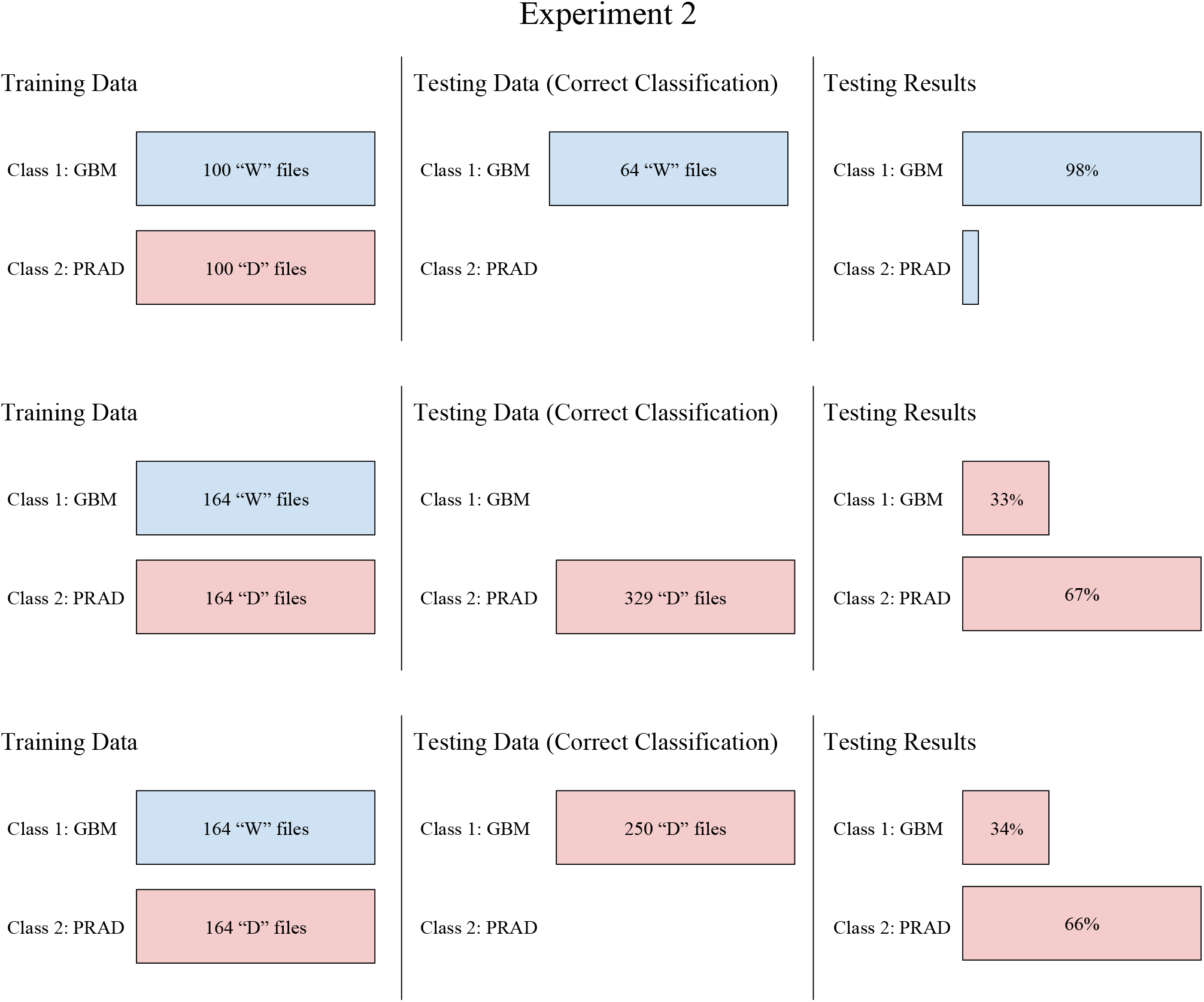
An illustration of experiment 2. Again, the placement of bars represent the labeling of the classes, which this time is separated by cancer type (GBM vs PRAD). We also continue our convention of coloring the hidden variable “D” type files red and “W” type files blue. Here, a classifier trained on cancers which differ in their file type appears to be successful in the first two test sets i.e., GBM “W” files and PRAD “D” files. The testing results for the third test set GBM “D” files, however, shows that the classification of GBM “D” files is very similar to that of PRAD “D” files. Hence, our machine learning algorithm has mistaken the D/W signal (blue vs red) for the cancer-type.

In Experiment 1, we consider the normal DNA of 414 patients with Glioblastoma Multiforme (TCGA-GBM). Of the 414 patients in TCGA-GBM, 164 patients had BAM files with analyte code “W” and the rest (250) had analyte code “D”. We train pairwise classifiers using gradient boosting^10,11^ (xgboost) to distinguish the “D” and “W” file types. At test time, 71% of the “D” files in the test set were correctly classified as “D” and 86% of the “W” files were correctly classified as “W”.

Strong differences, that contribute to this noise signature, are present in specific locations of the genome. Figure 3 shows the distribution of copy numbers at two different locations in “D” and “W” files in the normal DNA of TCGA-GBM patients. These differences, alongside point mutations, are present at many other locations and constitute the data amplification bias which is picked up by the xgboost classifier.

**Figure 3.**
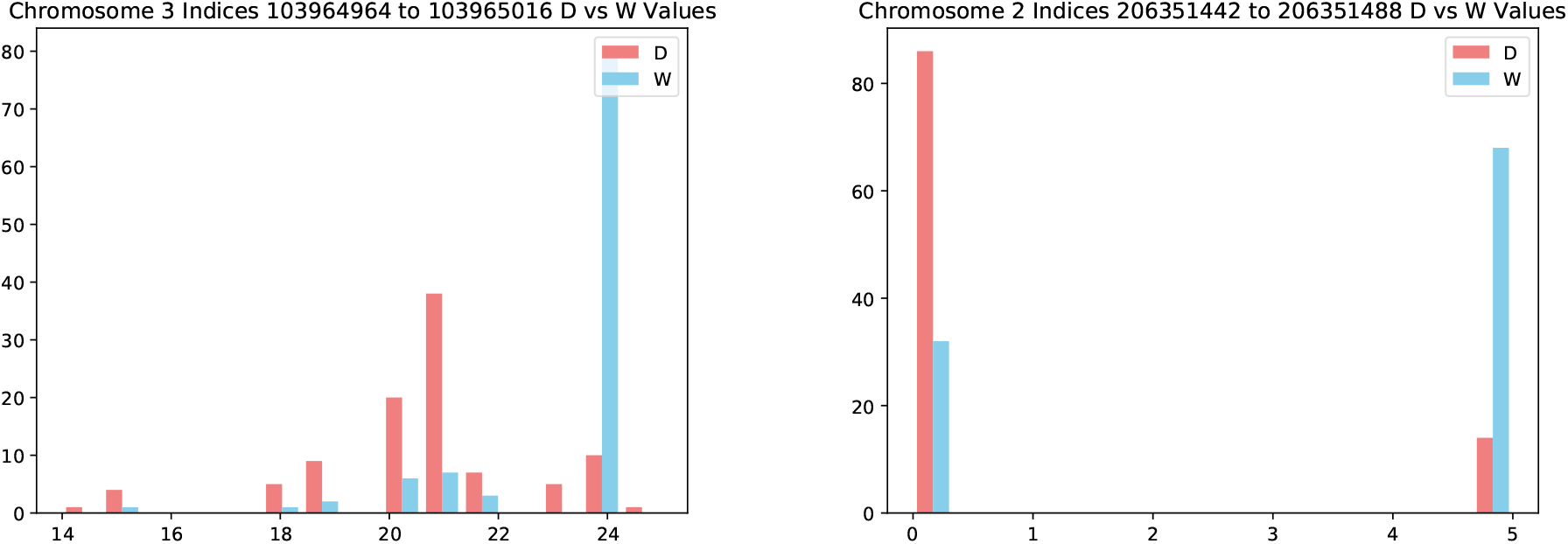
Left: The distribution of copy number at *chr3*: 103, 964, 964 – 103, 965, 016. It can be seen that for “W” files the distribution is concentrated at 24, however for D files the distribution is more spread out. Right: The distribution of copy number at *chr2*: 206, 351, 442 – 206, 351, 488. It can be seen that for most D files, the copy number is 0 showing the absence of the region, but the distribution of copy number for more than 60 W files is concentrated at 5.

In Experiment 2, we investigate the effect of this amplification bias on cancer prediction. For this purpose, we took 414 patients of TCGA-GBM (164 of “W” file type and 250 of “D” file type), and 493 patients from TCGA-PRAD, all with “D” file types. We build gradient boosting-based classifiers to distinguish between normal DNA of the GBM and PRAD cancer patients. For training, we used equal numbers of GBM type “W” and PRAD type “D” files.

As shown in Figure 2, GBM type “W” files comprise the test set and 98% are classified correctly as GBM files. When testing on PRAD type “D” files, the majority are also correctly classified as PRAD files. However, in the last part of the experiment, only 34% of TCGA-GBM type “D” files are correctly classified as TCGA-GBM, while the majority are incorrectly classified as TCGA-PRAD.

The results of experiment 2 (Fig. 2) show that files are classified as the correct cancer type *if* they match the file type of that cancer used in training. However, if the file type is that of the opposite cancer, they are misclassified, indicating that the amplification bias is stronger than any potential cancer signal. In fact, we get nearly identical classification results whether we use PRAD type “D” or GBM type “D” test files, implying that the cancer signal either does not exist or has been severely obscured. With regard to the question of cancer signal, we have separately analyzed “D” and “W” samples from TCGA in^7^. In light of these results, we discuss the repercussions of not considering amplification technique as a confounding factor in genomic studies.

## Discussion

As artificial intelligence is increasingly relied upon to substitute human reasoning, data bias is an acute concern. In particular, genomic data is highly susceptible to the source and technology by which it was generated, and hence may lead to false associations. Figure 1 establishes that there is a sufficient statistical difference between “D” and “W” file types for the xgboost classifier to identify the amplification bias present in the training set and distinguish file types. Figure 2 shows the effect this amplification bias has on the results of experiment 2 which was conducted to find cancer associations. By observing the test accuracies of 98% and 66% respectively, when testing was only done on GBM “W” file types or PRAD “D” file types, one might conclude that STR regions in normal DNA of GBM and PRAD patients have information about the 2 cancer types. However, when testing was done on GBM patients with “D” file types, we see contradictory results showing a stronger association of TCGA-GBM patients in the test set with TCGA-PRAD patients in the training set.

This contradictory result is observed due to the amplification bias. Since the training was done on TCGA-GBM patients with “W” file types and TCGA-PRAD patients with “D” file types, the hidden “D” vs. “W” signal in the training data overpowers the “GBM vs. PRAD” signal giving misleading classification accuracies for all test sets considered in experiment 2.

To account for these findings, prior to using any data-driven tool for finding phenotypical associations, a pre-check must be conducted in order to detect hidden variables, or “noise”, that might subside the true signal of interest. In our experiments, this hidden variable turned out to be the “D” and “W” file types. One potential approach to discover these hidden variables is *unsupervised learning*, where clustering techniques are used to find unknown associations in the data. Further approaches developed in *fairness in machine learning* literature can also be availed^2^.

## Methods

The classifiers were built using gradient boosting (xgboost) algorithm in the above experiments. 4-fold cross validation was done before building the final classifiers to identify the parameters. Short tandem repeats (pattern length ≤ 10) were extracted out of the WXS DNA and the number of copies (*d*) and the number of point mutations (*m*) in each of those tandem repeat regions are used as features (see Figure 4). Hence, for each tandem repeat region *i* in a genome, we compute *m_i_* and *d_i_*. If there are *N* tandem repeat regions in a DNA, the vector of 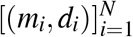 is called the mutation profile. Hence, each individual’s DNA is represented by this mutation profile, which serves as the input for building the gradient boosting based classifier. Samtools^9^ was used to process the BAM files. The repeat regions from the genome were extracted using Benson Tandem Repeat finder algorithm^3^ and the *m_i_* and *d_i_* values were estimated using the single block duplication history algorithm in Tang et al.^18^. Further details on the data availability and the Methods details are provided in^7^. The code for the pipeline used is available at http://paradise.caltech.edu/~sidjain/Codes.tar.gz.

**Figure 4.**
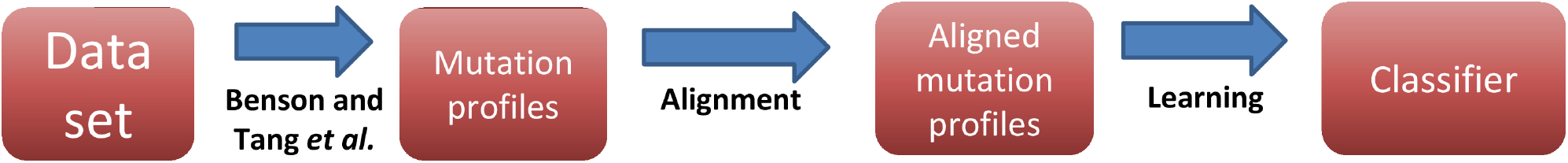
Pipeline to build a classifier for a given set of classes.

## Acknowledgements

This work was supported in part by The Caltech Mead New Adventure Fund and a Caltech CI2 Fund. The authors would like to thank Eytan Ruppin for his valuable advice and feedback.

## Ethics

The ethics approval to the TCGA data was granted by Caltech Institutional Review Board.

